# Forecasting fish recruitment using machine learning methods: A case study of arabesque greenling

**DOI:** 10.1101/2023.06.02.543365

**Authors:** Hiroshi Okamura, Shoko Morita, Hiroshi Kuroda

**Affiliations:** Japan Fisheries and Education Research Agency; Japan Fisheries Research and Education Agency

**Keywords:** Arabesque greenling, gradient boosting, machine learning, fish recruitment prediction

## Abstract

Fish recruitment prediction is a central topic in fisheries science. We use a machine learning method for predicting the recruitment of Arabesque greenling in Hokkaido, Japan. Biological, fisheries-related, and environmental factors were included in the predictive models as feature variables. A gradient boosting model (GBM) showed better predictive performance compared with a linear regression model (LRM) and a random forest model (RFM) in terms of relative bias and relative root mean square error for recruitment prediction in the last 5 years. The most influential feature for GBM was spawning stock biomass in the last year, followed by the fishing rate for older fish. The sea surface temperature (SST) was a very weak predictor in the GBM but was the most influential feature in the LRM. The difference among models suggests the importance of nonlinearity and variable interactions. Machine learning methods will greatly contribute to fish recruitment forecast and thereby sustainable fisheries management.

## Introduction

Fish recruitment is a piece of fundamental information for fisheries management and often shows dramatic variation. Accurate predictions of fish recruitment can contribute substantially to efficient fisheries management by increasing catch quota when recruitment will be high and decreasing catch quota when recruitment will be low, while maintaining sufficient spawning stock biomass. However, predicting fish recruitment is notoriously difficult. Fish recruitment is affected by many factors. In particular, larval survival is strongly dependent on environmental factors (Walters and Martell 2004). Szuwalski et al. (2015) reported that recruitment is more closely related to environmental factors than is spawning biomass over observed stock sizes for many stocks. In somewhat contrast, Myers (1998) stated that “The rarity of the use of environment-recruitment correlations is clear evidence against their general usefulness in assessments. Even if an environmental variable is important, it does not mean that it is key to the management of the fishery.” Recruitment prediction and identification of the underlying determinants are still key issues in fisheries science.

Arabesque greenling (*Pleurogrammus azonus*) is an important food fish in Japan and is often served and eaten in Japanese-style taverns. However, the catches, relative population size (catch per unit effort: CPUE), and estimated population sizes have declined, and a possible cause of these trends have been attributed to overfishing (Morita 2015, Morita 2017, Okamura et al. 2018). Increase of sea surface temperature (SST) due to global warming is another potential cause of the population decline because arabesque greenling prefer cold waters. Since 2012, stock management measures have been taken to ensure spawning stock by voluntarily regulating catch and fishing effort in response to declining population sizes (Hoshino et al. 2017). As a result of these stock management measures, the population size and spawning stock biomass in 2019 is on a recovery trend, and the catch has increased to 30,000 tons (Itaya et al. 2020). On the other hand, the population size of arabesque greenling is currently small compared to early 2000’s population size and the central component of catch is still young fish. Prediction of recruitment is therefore often more important than other things and the center of interest for arabesque greenling in Hokkaido.

Fish recruitment prediction has some difficulties: 1) the sample size is usually small (several tens to a hundred, at most) because population size estimation is annual, 2) multiple factors and higher-order interactions can influence predictions, and 3) big noise in fish recruitment data tends to mask important factors. The need to construct a complex model with many parameters, despite a small sample size, limits the use of conventional statistical models such as a linear regression. Fortunately, for cases in which the sample size is smaller than the number of parameters, new statistical approaches are now available, including machine learning methods (James et al. 2013).

In the present study, we predict recruitment of the northern Hokkaido (NH) arabesque greenling stock using machine learning methods including factors related to fish biology, fisheries, and environmental conditions. Machine learning methods improved the accuracy and precision of fish recruitment prediction compared with a conventional linear regression model and clarified the factors that are most influential in terms of prediction. We discuss the limitations of conventional models and merits of machine learning methods based on the application of these approaches to arabesque greenling.

## Materials and Methods

### Stock size and fishing activities

The population sizes and fishing rates for the NH arabesque greenling stock were estimated on a half-year basis using the tuned virtual population analysis (VPA) in the annual stock assessment (Morita et al. 2022a). In this study, the spawning stock biomass (*SSB*) is defined as

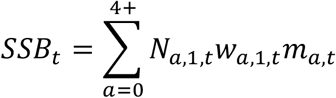

where *N*_*a,s,t*_ is the number of fish at age *a* (*a* = 0, …, 4+) in season *s* (*s* = 1 (1st half) or 2 (2nd half)) of year *t* (*t* = 1985, …, 2021), *w*_*a*,s,*t*_ is the mean weight at age *a* in season *s* of year *t*, and *m*_*a,t*_ is the maturity proportion at age *a* of year *t*. The corresponding recruitment (*R*_*t*_) in year *t*, which is the target (or response) variable to be predicted in this study, is the number of fish at age 0 and season 1 (i.e., *R*_*t*_ = *N*_0,1,*t*_).

The connection between *R* and *SSB*, called the stock-recruitment (SR) relationship, which gives fundamental information for estimating the maximum sustainable yield (MSY) (e.g., Ichinokawa et al. 2017, Okamura et al. 2021). The hockey-stick type SR relationship was used for NH arabesque greenling (Morita et al. 2022b). *R* is also affected by fishing activities and environmental changes directly and indirectly. We therefore adopt *SSB*, the fishing rate (*U*), *SST*, and past *R* (i.e., autoregression) as potential factors influencing the prediction of recruitment.

### Sea surface temperature

Because survival rates of larval and juvenile fish are strongly affected by SST (Kuroda et al. 2020), we adopted SST as a potentially useful environmental indicator for predicting future fish recruitment. Mean SST is increasing annually because of global warming (Kuroda et al. 2020, Hayashi et al. 2022). Arabesque greenling use offshore for feeding ground (Kambara 1957) and inshore for spawning ground (Gomelyuk 1988, Munehara and Markevich 2003, Takashima et al. 2016). According to the reports on the range of temperature for spawning season by Miyaguchi (1983) and Sakaguchi et al. (2022), arabesque greenling prefer cold waters and are therefore expected to shift northward, possibly outside the fishing area, as mean SST becomes higher. The time series of mean SST from 1982 to 2020 was broken into four seasons (Winter: January to March, Spring: April to June, Summer: July to September, and Autumn: October to December) and two large areas (OK [Okhotsk Sea] and SJ [Sea of Japan]). Principal component analysis (PCA) was then conducted separately for each segment. The first and second principal components were used as predictors for future fish recruitment. SST for each segment was discriminated using the notation *SST*_*t,s,a,p*_ where *t* is the year (*t* = 1985, …, 2020), *s* is the season (*s* = Winter, Spring, Summer, and Autumn), *a* is the area (*a* = OK or SJ), and *p* is the *p*th principal component (PC: *p* = 1 or 2). The first PC generally corresponded to the trend in SST and the second PC corresponded to the standard deviation of SST, while the relationship between the third PC and higher-order statistics (skewness and kurtosis) was weak (Fig. S1 in Supplementary Materials A). Because the cumulative proportion of variance explained by the first and second PCs was generally greater than 90% and the interpretation of the third PC is difficult, only the first two PCs were used as covariates in our analysis.

### Machine learning method for predicting future recruitment

A gradient boosting model (GBM) was used for forecasting fish recruitment. GBM uses a combination of many decision trees to improve the prediction skill. A predictive model *f*(*x*) is constructed by minimizing the loss function *L*(*y, f*(*x*)), where *y* is the response variable (fish recruitment) and *f*(*x*) is a nonlinear function of feature vector (covariates) *x* (SSB, SST, etc.). In GBM, the predictive model *f*(*x*) is obtained sequentially as follows (Ridgeway 2020):

1. For *t* = 0, assume that *f*_0_(x) is a constant *ρ* by minimizing 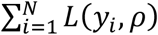 and set *t* = 1.
2. Then calculate the negative gradient 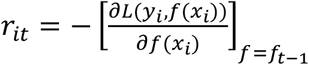 and fit the regression tree *g*(*x*_*i*_) to predict *r*_*it*_.
3. Calculate 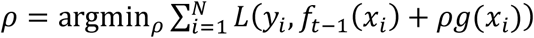.
4. Update *f*_*t*_(*x*) = *f*_*t*-1_(*x*) + *ρg*(*x*) and iterate 2 to 4 for *t* = 1, …, *T*.

The use of the negative gradient in step 2 above stems from the gradient descent method, an optimization algorithm to find the minimizer of the loss function *L*(*y*). In the gradient descent method, *y* is updated by *y*_*t*+1_ = *y*_*t*_ − *yL*′(*y*_*t*_) (*γ* is a small number) such that *y* gets the minimizer of *L*(*y*). Therefore, learning of the negative gradient corresponds to learning of the residual *y*_*t*+1_ − *y*_*t*_. That is to say, GBM sequentially learns the residuals that have not been learned in the previous steps. Even if a learner for each residual is weak, a final ensemble of weak learners becomes a powerful predictive machine.

To predict fish recruitment, we define log(*R*_*t*_) as a response variable. The feature vector is then given by

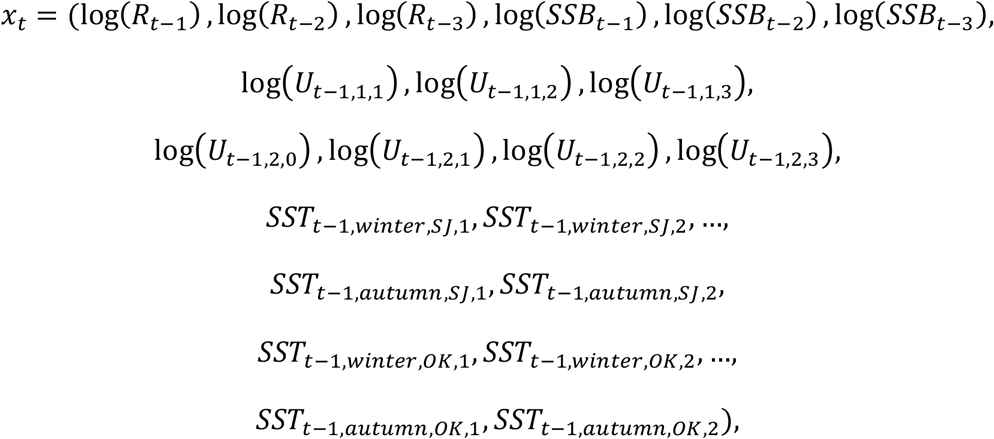

where *R*_*t*_ is the recruitment in year *t, SSB*_*t*_ is the spawning stock biomass in year *t, U*_*t*,1,*a*_ is the fishing rate at age *a* in the first half of year *t, U*_*t*,2,*a*_ is the fishing rate at age *a* in the second half of year *t*, and *SST*_*t,s,z,p*_ is the *p*th principal component of mean sea surface temperature predicted by PCA in year *t*, season *s*, and area *z*. To examine the influence of time lags, *R* and *SSB* of the past 3 years were taken into account, whereas *U* and *SST* of the past 1 year were used because the influence of these variables is propagated into *R* and *SSB*. The starting year of the response variable, log(*R*_*t*_), is therefore 1988 (1988 – 3 = 1985 is the first year of available population size and fishing rate time series). The fishing rate *U*_*t*-1,1,0_ is excluded from the vector of feature variables because it is mostly zero.

We use the R package “gbm” (Ridgeway 2020) for the gradient boosting regression. Five hyperparameters in gbm were tuned to maximize performance: n.trees (total number of fitted trees, *T*), bag.fraction (proportion of the training dataset randomly selected in each step of learning, where values of <1 make GBM stochastic), interaction.depth (maximum depth of each tree corresponding to the maximum level of interactions allowed), shrinkage (learning rate, where small values correspond to slow convergence, requiring a large number of n.trees), and n.minobsinnode (minimum number of observations in the terminal nodes of trees). We used a grid search to find the best set of hyperparameter combinations. The grid was set to the following for each hyperparameter: bag.fraction ∈ {0.80, 0.85, 0.90, 0.95, 1.00} (by 0.05), interaction.depth ∈ {1, 2, …, 8}, shrinkage ∈ {0.200, 0.225, …, 0.500} (by 0.025), and n.minobsinnode ∈ {2, 3, …, 8}. n.trees was commonly set to 5,000 for each of combination of other hyperparameters, and the optimal n.tree less than or equal to 5,000 was selected by leave-one-out cross validation (loocv) based on the Gaussian distribution (i.e., the squared error function). The set of hyperparameters with the minimum loocv over all combinations of hyperparameters was finally selected.

The dataset was divided into training and test datasets. The training dataset included data from 1988 to 2016 and the test dataset was from 2017 to 2021. The model with the best set of hyperparameters based on the training dataset was used for predicting the test dataset. The prediction of recruitment in year *t* (*t* = 2017, .., 2021) for the test dataset was conducted using the dataset from 1988 to year *t* – 1 (i.e., one-year ahead prediction). The prediction performance was evaluated by the relative mean absolute bias (RMAB)

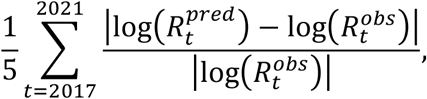

and the relative root mean square error (RRMSE) of log-recruitments:

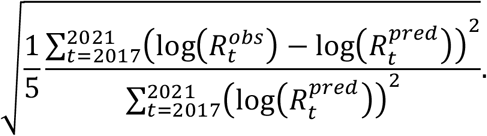

The impact of feature variables on recruitment prediction was evaluated by a variable importance plot (VIP) and the relationships between the feature variables and the recruitment were visualized by a partial dependence plot (PDP) (James et al. 2013). The VIP was based on the reduction of squared errors by the elimination of each variable and consequently each variable was ranked in order based on the magnitude of squared error reduction. The PDP was calculated by changing the target variable over the variable of interest with the other variables averaged over their observed ranges.

For comparison, the simple linear regression model (LRM) and random forest model (RFM) were fitted to the same training dataset (For LRM, however, only the 1st PCs as SST were included and the 2nd PCs were excluded because of convergence problems) and log-recruitment was predicted using the same test dataset (Supplementary Materials

B). Model selection based on AIC was used for LRM and recruitment was predicted using the best model. The same framework used for the above GBM was used for RFM without tuning any hyperparameters because RFM is robust for the selection of hyperparameters (“mtry” (the number of variables used for each split of the tree regression), an important hyperparameter for RFM, is set to one-third of the number of feature variables as the default setting (James et al. 2013)).

## Results

The best set of hyperparameters for GBM selected by cross validation was interaction.depth = 3, shrinkage = 0.300, n.minobsinnode = 5, bags.fraction = 0.80, and n.trees = 9. We refitted the predictive model with the best set of hyperparameters to the whole dataset (i.e., the training data + test data) and identified this as the best GBM. The VIP for the best GBM showed that SSB is the most important predictor, followed by the fishing rate at age 2 in the last half of the year (*u*_22_) (Fig. 1). Next, the fishing rates at age 3 and SST in winter at Okhotsk emerged, though their effects were very weak. The recruitment of the last year (*R*_1_) was the sixth most important variable for recruitment prediction, although its impact was much smaller than those of the predictors with higher ranks. This effect of *R*_1_ implies that there may be environmental changes other than SST with weak effects on recruitment prediction.

**Figure 1.**
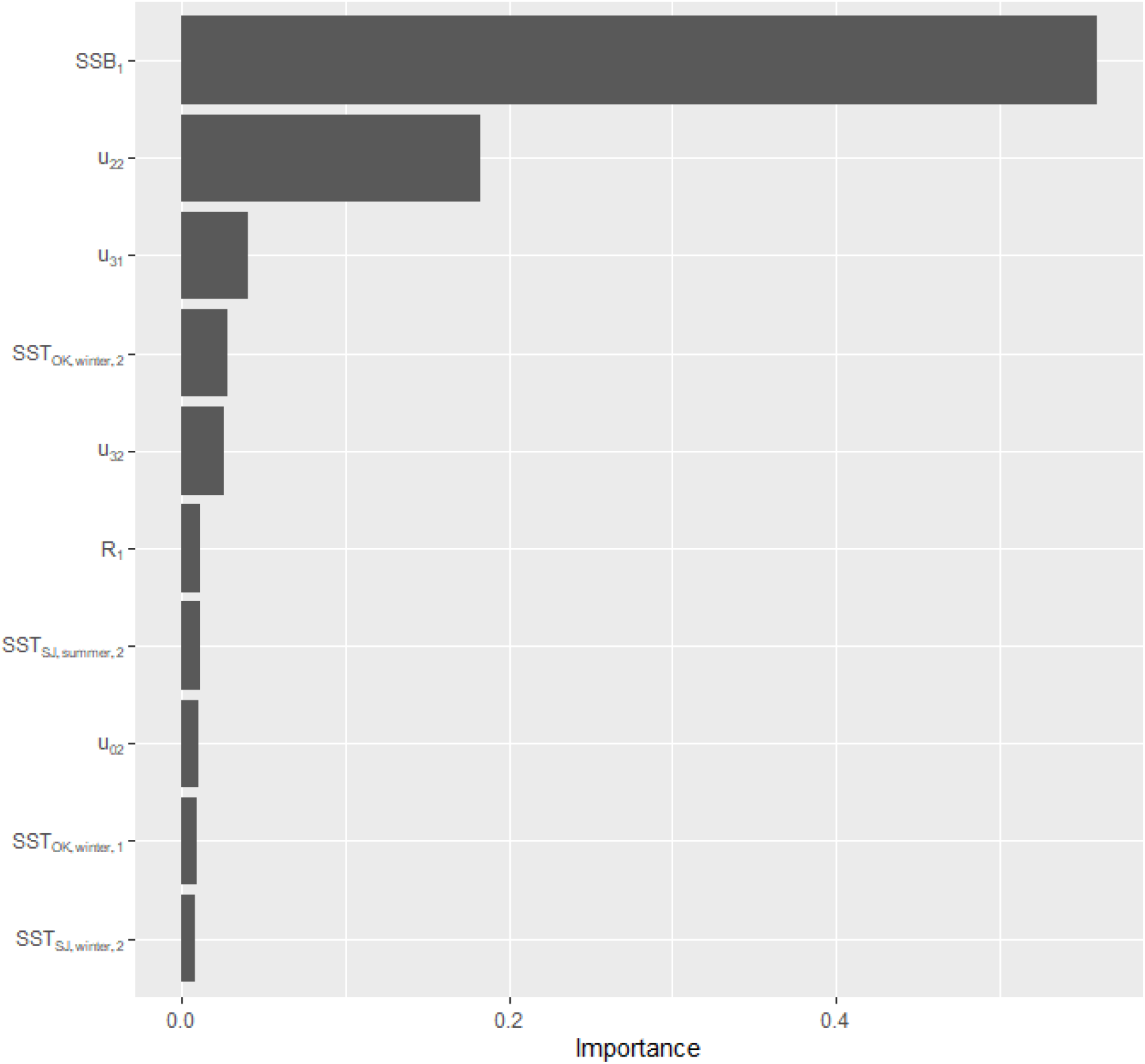
Variable importance plots for GBM. SSB_1_: SSB in the last year, u_22_: fishing rate at age 2 and season 2, u_31_: fishing rate at age 3 and season 1, SST_OK,winter,2_: 2nd PC of SST at Okhotsk in winter, u_32_: fishing rate at age 3 and season 1, R_1_: recruitment in the last year, SST_SJ,summer,2_: 2nd PC of SST at Sea of Japan in summer, u_02_: fishing rate at age 0 and season 2, SST_OK,winter,1_: 1st PC of SST at Okhotsk in winter, SST_SJ,winter,2_: 2nd PC of SST at Sea of Japan in winter.

The PDPs for the ten most significant predictors generally depicted simple step functions (Fig. 2). SSB was related to increased recruitment in the next year when it exceeds 40.7 thousand tons, the mean value over the range of SSB corresponding to dramatic recruitment changes. The fishing rate *u*_22_ predicted lower recruitment when it exceeded 0.32, the mean value over the range of *u*_22_ corresponding to dramatic recruitment changes. Although the relationships between remaining eight predictors and recruitment changes were very weak, the general trends of some variables were consistent with our expectations. For example, recruitment in the target year had a positive correlation with the recruitment in the last year (*R*_1_) and negative correlations with the fishing rate at age 3 (*u*_32_) and SST_OK,winter,1_. However, the trends in some variables were difficult to interpret (for example, the fishing rate at age 0 (*u*_02_) had a negative correlation with recruitment).

**Figure 2.**
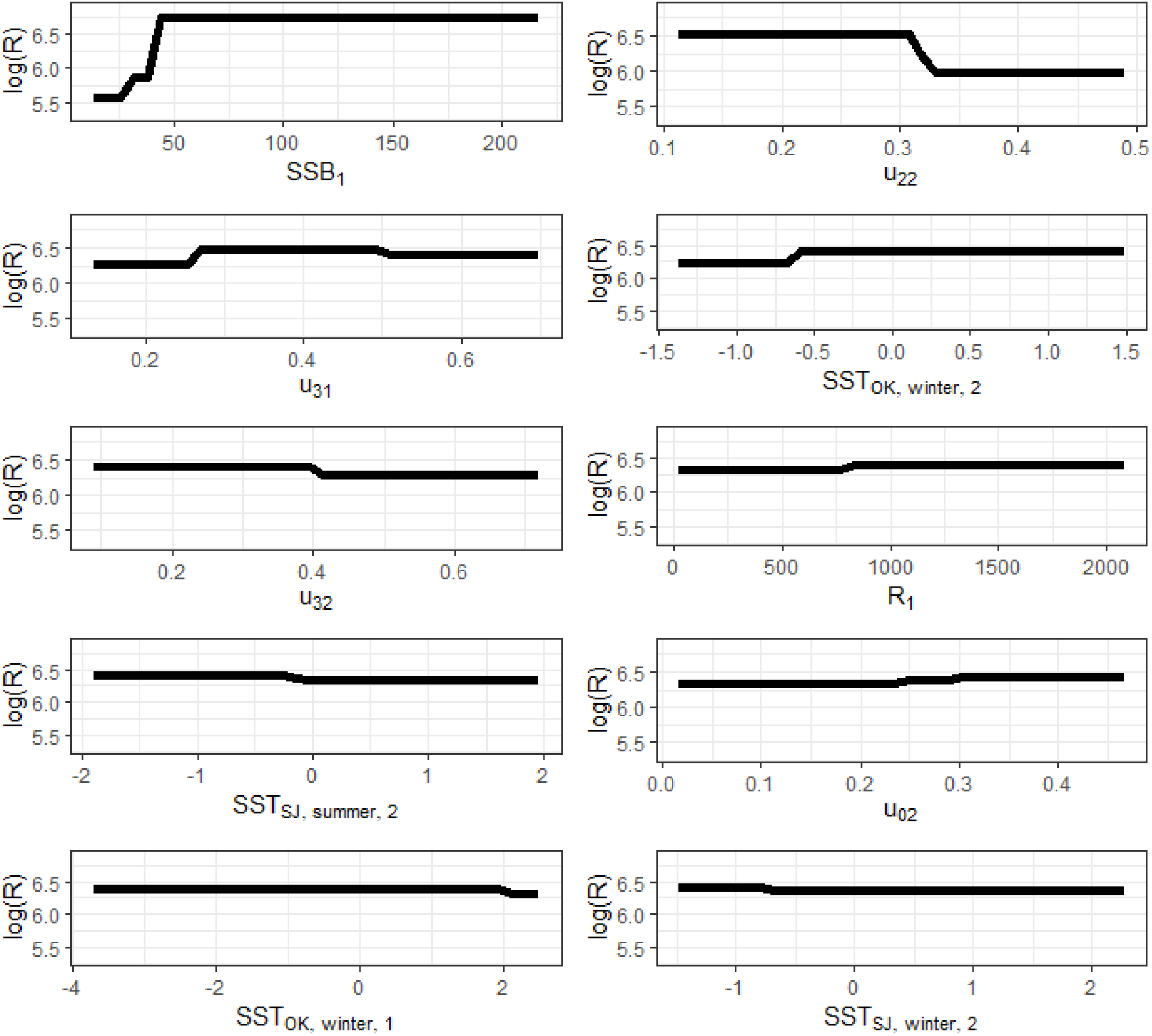
Partial dependence plots for GBM. The variable names on the horizontal axis are the same as those of Fig. 1.

The predicted recruitments were consistent with the observed values (Fig. 3). The correlation coefficient between the observed and predicted recruitments for GBM was 0.94, whereas those for LRM and RFM were 0.78 and 0.76, respectively. RMAB for GBM was 0.066, whereas those for LRM and RFM were 0.122 and 0.075, respectively. RRMSE for GBM was 0.036, whereas those for LRM and RFM were 0.064 and 0.037, respectively. Recruitment predicted from 2017 to 2021 using the dataset up to the target year showed different results depending on the prediction model (Fig. 4). GBM traced the variation in observed recruitment well, although the deviation in 2019 was relatively large. LRM failed to effectively predict both the trend and variation, and the RFM was nearly linear, ignoring the variation around the trend as the errors. The influential features in RFM were similar to those of GBM, such as *SSB* and *u*_22_, whereas LRM indicated that summer and autumn SSTs and fishing rates at age 3 as important predictors that were substantially different from the important variables in the other two models (Fig. S2 in Supplementary Material B).

**Figure 3.**
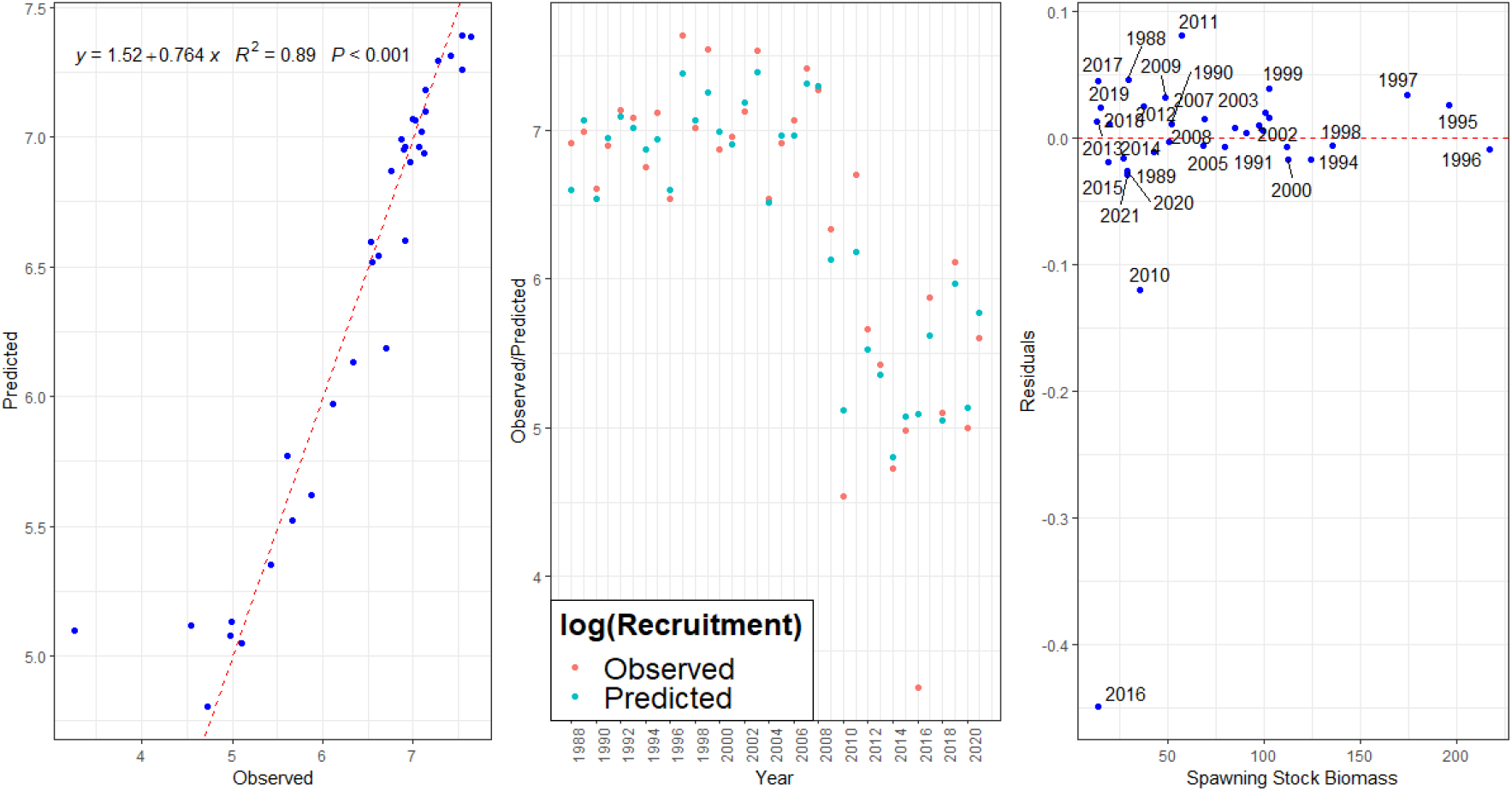
Goodness of fit for GBM. Left: comparison plot for observed vs. predicted where the red broken line is the 1:1 line, Middle: year-based plot for observed vs. predicted, and Right: residual plot for observed minus predicted.

**Figure 4.**
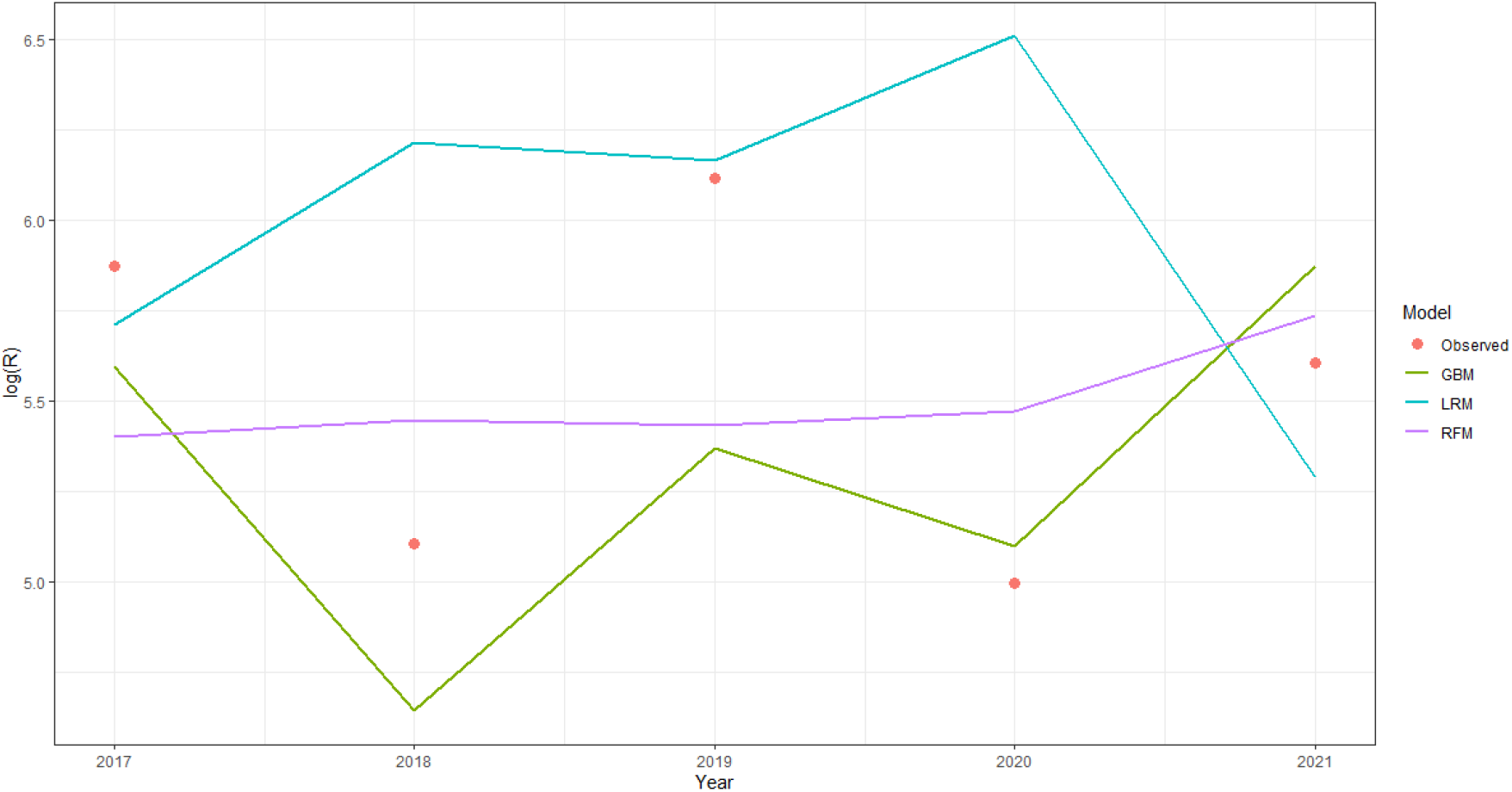
Prediction for 2017–2021 recruitments using GBM, LRM, and RFM. Red dots indicate the observed log recruitments.

## Discussion

GBM effectively predicted recruitment in the target year, although it sometimes failed to give accurate estimates (e.g., in 2016 and 2019) (Figs. 3 and 4). LRM showed considerably worse performance compared with those of the machine learning methods (i.e., GBM and RFM) and was not effective for predicting future recruitment (Fig. 4). The most important feature in both GBM and RFM was SSB in the last year, while the most important feature in LRM was the 1st PC of summer SST in the Sea of Japan. In LRM, SSB had little or no effect on the prediction of recruitment. The linear functional form in LM might not have captured the important features, as corroborated by the fact that the relationship between SSB and R in GBM was a nonlinear step function (Fig. 2). Although we can extend LRM to more advanced models, including GLM, GLMM, GAM, and GAMM, to deal with nonlinearity and complex model structure (Zurr, Ieno, and Smith 2007, Wood 2017), LRM and its extensions cannot handle parameter counts exceeding the sample size. This limits the ability to identify important features for prediction. However, tree-based machine learning methods, including GBM and RFM, can generally deal with such nonstandard situations naturally because feature counts more than sample size do not hinder the model construction (James et al. 2013).

GBM was superior to RFM and LRM in terms of fitting and RMAB, and outperformed LRM and was nearly equivalent to RFM in terms of RRMSE. Although RFM was comparable to GBM in terms of RRMSE, RFM failed to capture the variation in recruitment and predicted a nearly linear trend (Fig. 4), probably due to lack of fit to past data. Because GBM was best in terms of relative bias (RMAB) and its performance on RRMSE was almost equivalent to that of RFM, we recommend using GBM for future recruitment prediction of the NH arabesque greenling stock.

The most influential feature in GBM and RFM was SSB in the last year (Fig. 1 and Fig. S2). The fishing rate at age 2 was also influential in both GBM and RFM. Not to mention the effect of SSB, the observation that low fishing pressure for higher ages predicted higher recruitment suggests the importance of conserving adult populations. Although some effects of SST-related features were observed, these effects were generally weaker than those of SSB and fishing rates at higher ages. In LRM, the summer SST in the Sea of Japan was the most important feature. Because this SST is related to the general trend in SST, this matches the reported preference of arabesque greenling for cool waters and the northward shift of the species to avoid the influx of warm water due to global warming (Miyaguchi 1983, Sakaguchi et al. 2022). However, considering the goodness of fit and the predictive performance of LRM, the predictive value of SST would be much lower and the difference between a conventional regression method (LRM) and machine learning methods (GBM and RFM) emphasizes the importance of accounting for nonlinearity and many features for fish recruitment prediction. In GBM, the effect of SST was weak and, if any, the variation was more effective than the trend.

The relationship between R and SSB for GBM was consistent with the hockey-stick stock-recruitment function (Fig. 2), the stock-recruitment function used in the actual stock assessment (Morita et al. 2002b). The transition point from low to high recruitment was about 40 thousand tons and was close to the limit reference point (SB_60%MSY_: spawning stock biomass that produces catch at equilibrium equal to 60% of MSY), 30 thousand tons, estimated by the stock assessment in 2022 (Morita et al. 2022b). Keeping SSB high and reducing fishing pressure for older individuals can contribute to maintain higher recruitment over a long time. The prediction performance for recruitment in 2016 was substantially bad in comparison with results in other years (Fig. 3), but the reason is not obvious at present. Such extreme population decline might be due to some environmental factors not included in the model and/or some biological factors such as the Allee effect (Hutchings 2021).

Features useful for prediction are not necessarily the causes of high or low recruitment because of the difference between prediction and causal relationship (McElreath 2020). Therefore, the relationship between R and SSB revealed by GBM does not necessarily reflect a causal relationship. However, an understanding of causal relationships is not absolutely necessary for sustainable fisheries management. The predictive value of SSB for future recruitment by machine learning methods supports the reliability of the current management procedure based on the functional relationship between R and SSB.

GBM improved recruitment prediction substantially over that of conventional regression methods. Machine learning methods are thus promising for improving fish stock assessment and management. For the NH arabesque greenling stock, biology-related and fisheries-related information is more important than environmental information for future recruitment prediction. However, some fishes, such as pelagic fishes, would be more sensitive to environmental factors. The application of flexible approaches, such as machine learning methods, can clarify the important factors for the prediction of fish recruitment and contribute to sustainable fisheries management in the future.

