## Supplemental materials for "Forecasting fish recruitment using machine learning methods: A case study of arabesque greenling"

Forecasting fish recruitment using machine learning methods: A case study of arabesque greenling
Supplementary Materials (Online Only)

Hiroshi Okamura, Shoko Morita, and Hiroshi Kuroda

**Supplementary Materials A**: Principal component analysis of sea surface temperature

Monthly sea surface temperature (SST) was extracted from NOAA NCEI GHRSST Level-4 OI-SST v2.1 (https://podaac.jpl.nasa.gov/dataset/AVHRR_OI-NCEI-L4-GLOB-v2.1, DOI: https://doi.org/10.5067/GHAAO-4BC21). We used SST for the 20 quadrats (Fig. S1) from 1982 to 2020. For this study, we further extracted 15 quadrats in Okhotsk Sea and Sea of Japan, which are within habitats the NH arabesque greenling stock, out of 20 quadrats. Since SSTs between quadrats have high correlations (the minimum correlation is 0.946), we first divided the SST time series into two large areas (OK[Okhotsk Sea] and SJ[Sea of Japan]) and four seasons (Winter: January to March, Spring: April to June, Summer: July to September, and Autumn: October to December). We conducted PCA for each divided mean SST time series using the R function "prcomp". The *p*th principal component (PC) was obtained using the "predict" function. The 1st and 2nd PCs generally explained more than 95% in the total variance.

To help interpretation about what the pth PC is, we compared the pth PC with some basic statistics (mean, standard deviation, skewness, and kurtosis). The 1st PC was consistent with the mean with high correlations and the 2nd PC was consistent with the standard deviation with high correlations (Fig. S2). However, the 3rd PC was not consistent with any specific statistics. We therefore used only the 1st and 2nd PCs for machine learning methods.

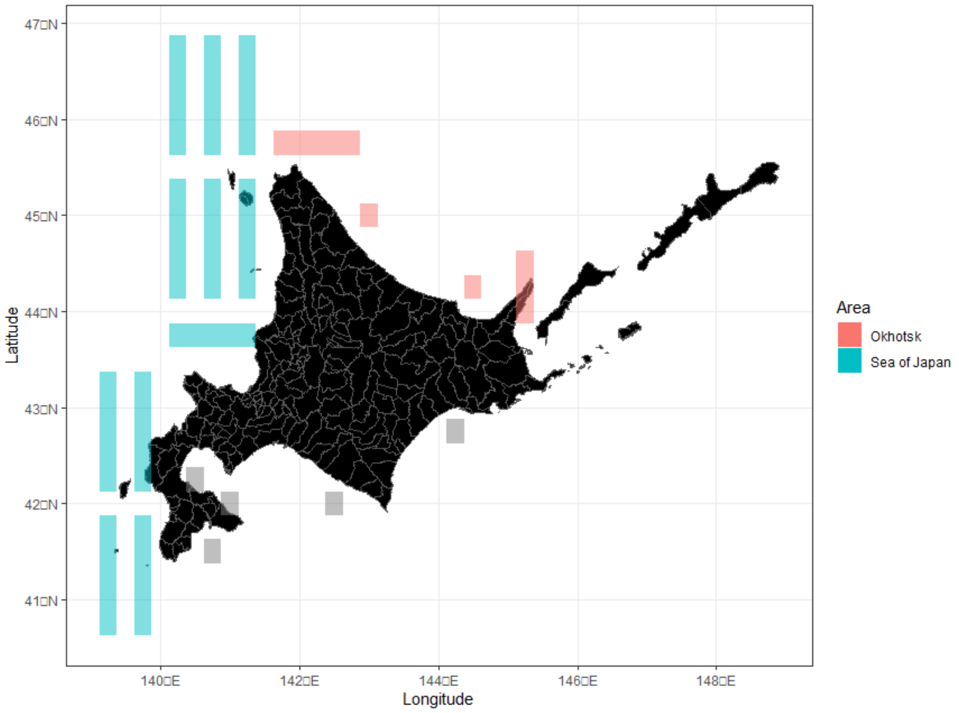


Figure S1. A map for SST sampling stations. Green quadrats correspond to Sea of Japan and red quadrats correspond to Okhotsk. Grey ones were just excluded before analysis.

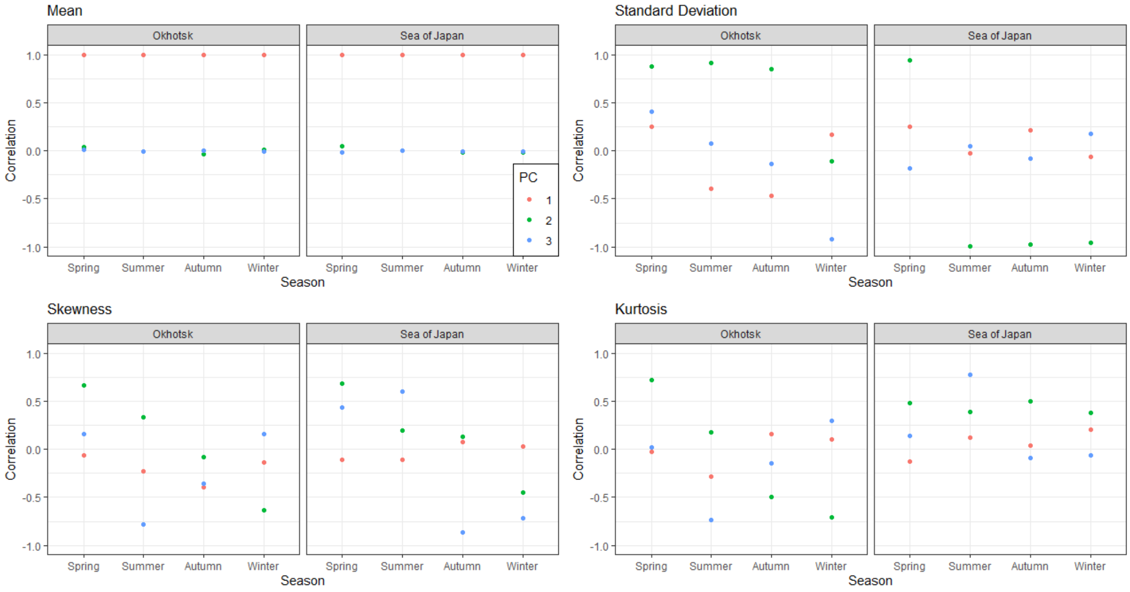


Figure S2. Comparison between the basic statistics (mean, standard deviation, skewness, and kurtosis) and the *p*th PC ($p$ = 1, 2, and 3).

**Supplementary Materials B**: Results Using Linear Regression Model and Random Forest Model

For comparison, we applied a simple linear regression model (LRM) and a random forest model (RFM) for the same dataset that was fitted by GBM. For LRM, the same feature vector was not used because parameter counts are too many. We used the following three feature vectors and adopted the lowest RMAB and RRMSE for LRM:

1) Only the 1st PC on SST and others same as the feature vector for GBM,

2) Only the 2nd PC on SST and others same as the feature vector for GBM, and

3) Only the 1st PC on SST, log(SSB) instead of SSB, log(R) instead of R, and others same as the feature vector for GBM.

The results of RMAB and RRMSE are provided in Table S1. The first LRM had the smallest RMAB and RRMSE. However, the third LRM had the smallest AIC. The 1st and 2nd LRMs had SSTs as important variables, whereas the 3rd LRM had SSB in the last year as the most important variable and any SST variable was excluded from the AIC best model. This suggests the importance of nonlinearity, the effect of SST depends on a specific model structure and seems dubious.

The variable importance plots (VIP) for the RFM and the best LRM in terms of RMAB and RRMSE are shown in Fig. S3. Whereas LRM had SSTs as important variables, the important variables in RFM were generally similar to those in GBM (Fig. 1).

Table S1. RMAB and RRMSE for the three LRMs.

| LRM | RMAB | RRMSE |
|---|---|---|
| 1 | 0.1225 | 0.0637 |
| 2 | 0.1290 | 0.0676 |
| 3 | 0.1479 | 0.0863 |

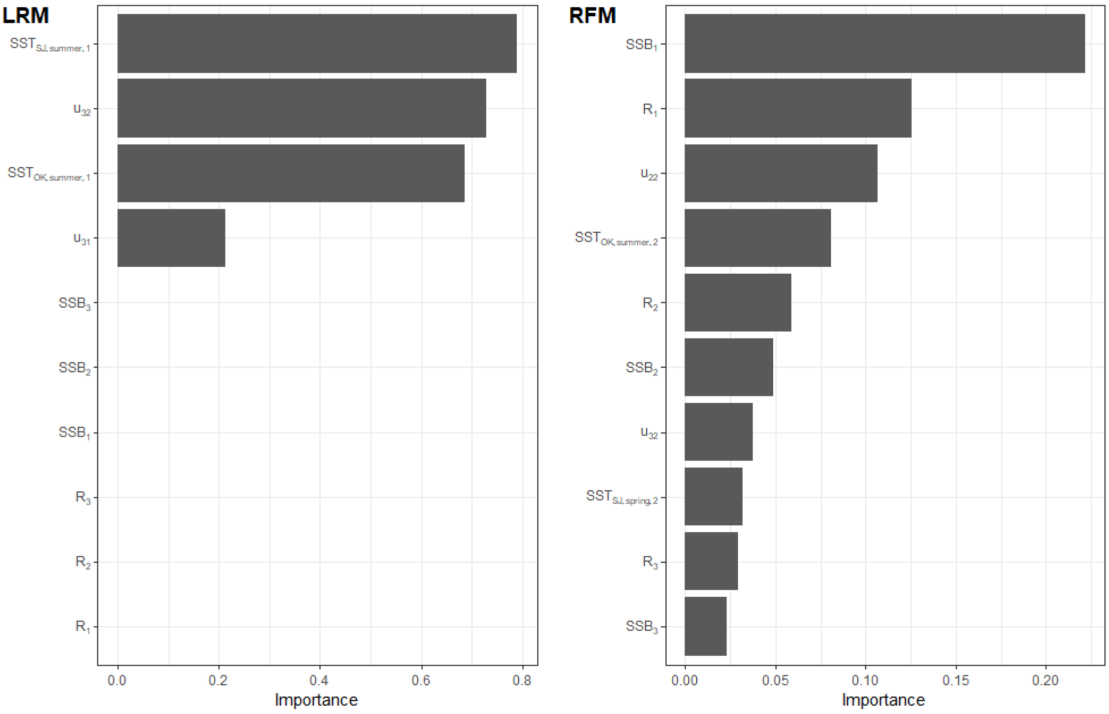


Figure S3. Variable importance plots for LRM (Left) and RFM (Right). $SSB_t$: SSB before *t* years ($t$ = 1, 2, and 3), $R_t$: R before *t* years ($t$ = 1, 2, and 3), $u_{as}$: fishing rate at age *a* and season *s*, and $SST_{Area, Season, p}$: *p*th PC of SST at *Area* (OK or SJ) in *Season* (winter, spring, summer, and autumn).
